# Synaptic Expression of TAR-DNA-Binding Protein 43 in the Mouse Spinal Cord Determined Using Super-Resolution Microscopy

**DOI:** 10.1101/2022.08.29.505610

**Authors:** Matthew J. Broadhead, Katherine Doucet, Owen Kantelberg, Fei Zhu, Seth GN Grant, Mathew H Horrocks, Gareth B. Miles

**Author notes:** First and Corresponding Author;.

## Abstract

Cellular inclusions of hyperphosphorylated TAR-DNA-Binding Protein 43 (TDP-43) are a key hallmark of neurodegenerative diseases such as Amyotrophic Lateral Sclerosis (ALS). ALS is characterised by a loss of motor neurons in the brain and spinal cord that is preceded by early-stage changes in synaptic function that may be associated with TDP-43 pathology. However, there has been little characterisation of the synaptic expression of TDP-43 in spinal cord synapses. This study utilises a range of high-resolution and super-resolution microscopy techniques with immunolabelling, as well as an aptamer-based TDP-43 labelling strategy visualised with single-molecule localisation microscopy, to characterise and quantify the presence of phosphorylated TDP-43 (pTDP-43) in spinal cord synapses. We observe that TDP-43 is expressed in the majority of spinal cord synapses as nanoscale clusters as small as 60 nm in diameter. Synaptic TDP-43 expression is more frequently associated with presynaptic terminals than postsynaptic densities, and is more enriched in VGLUT1-associated synapses, compared to VGLUT2-associated synapses. Our nanoscopy techniques showed no difference in the subsynaptic expression of pTDP-43 in the ALS mouse model, SOD1^G93a^ compared to healthy controls. This research characterizes the basic synaptic expression of TDP-43 with nanoscale precision and provides a framework with which to investigate the potential relationship between TDP-43 pathology and synaptic pathology in neurodegenerative diseases.

## Introduction

Amyotrophic Lateral Sclerosis (ALS) is characterised by a progressive loss of motor control due to a loss of motor neurons (MNs) in the spinal cord, brain stem, and motor cortex. Prior to MN loss, structural and functional changes in synapses have been reported between neurons in both the brain and spinal cord (Bączyk et al., 2020; Broadhead et al., 2022; Fogarty et al., 2015, 2016; Jiang et al., 2019). Synaptopathy in ALS is characterized by early stage hyper-excitability, that may induce excitotoxicity, and a later stage loss of vulnerable synapse subtypes (Broadhead et al., 2022; Fogarty, 2019; Van Zundert et al., 2008). The molecular mechanisms of ALS synaptopathy, however, are not fully understood.

TAR-DNA-Binding Protein 43 (TDP-43) is a ubiquitously expressed DNA/RNA binding protein that provides critical roles in RNA splicing, translation and transport in the nucleus and cytoplasm of neurons and glia in the nervous system (Ayala et al., 2008, 2011; Tollervey et al., 2011). Mutations in the gene encoding TDP-43 (TARDBP) have been identified in 4-5% of familial ALS cases and 1% of sporadic ALS cases (Millecamps et al., 2010). The aberrant accumulation of misfolded, hyper-phosphorylated TDP-43 (pTDP-43) in the cytoplasm of neuronal and glial cells also occurs in over 90% of all ALS cases (Mackenzie et al., 2010).

TDP-43 may play a role at synapses in both healthy and diseased conditions (Ling, 2018). TDP-43-bound mRNAs encode proteins that have roles in presynaptic and postsynaptic function, such as the glutamate receptor GluA1, glial excitatory amino acid transporter-2 (EAAT2) and microtubule-associated protein Map1b (Colombrita et al., 2012; Godena et al., 2011; Sephton et al., 2011; Tollervey et al., 2011). The expression of TDP-43 has been observed in the dendrites of cultured rat hippocampal neurons and colocalised with markers of the postsynaptic density (PSD) of excitatory synapses (Wang et al., 2008). Genetic alteration of the expression levels of TDP-43 in ALS mouse models is associated with both increased and decreased dendritic branching and synapse number, suggesting a role in regulating the structural plasticity of synapses (Fogarty et al., 2016; Jiang et al., 2019; Ling, 2018; Majumder et al., 2012). Analyses from human patients with ALS and the related cognitive disorder, frontotemporal dementia (FTD), suggest a strong correlation between TDP-43 pathology, synapse loss and the severity of cognitive symptoms (Henstridge et al., 2018).

Taken together, synaptic dysfunction may be linked to TDP-43 pathology. However, there has been little characterisation of the synaptic expression of TDP-43 in spinal cord synapses. MNs, for example, receive numerous different types of synaptic inputs, from excitatory, inhibitory and modulatory synapses derived from descending inputs from the brain, local spinal cord interneurons and sensory neurons. Not all synapses are equal, as some may be more selectively targeted in certain disease conditions (Bączyk et al., 2020; Broadhead et al., 2022; Herron & Miles, 2012; Mentis et al., 2011; Sunico et al., 2011). Understanding the expression of TDP-43 in different types of synapses in the spinal cord may provide insights into the selective vulnerability of certain synapses and pathways in ALS and therefore advance knowledge of the pathogenic mechanisms underlying this devastating disease.

In this study, we have used high-resolution and super-resolution microscopy techniques to visualise and assess the presence of phosphorylated TDP-43 (pTDP-43) at synapses within the mouse spinal cord.

## Methods

### Animals and ethics

All procedures performed on animals were conducted in accordance with the UK Animals (Scientific Procedures) Act 1986 and were approved by the University of St Andrews Animal Welfare and Ethics Committee. The B6SJL-TgN(SOD1-G93A)1Gur/J (SOD1^G93a^) mouse line was kindly provided by Dr Richard Mead. PSD95-eGFP mice were originally obtained from Prof. Seth Grant (University of Edinburgh). The PSD95-eGFP mouse is a genetically engineered in-frame fusion knock-in mouse that fuses the enhanced green fluorescent protein (eGFP) to the C-terminal of the endogenous PSD95 protein, enabling the visualisation of postsynaptic densities (PSDs) of excitatory synapses throughout the nervous system. As characterised previously, SOD1^G93a^ x PSD95-eGFP progeny display similar weights and behavioural phenotypes as expected from ‘pure’ SOD1^G93a^ mice, with the onset of hind-limb tremors and reduced hind-limb splay by approximately 75 days of age (Broadhead et al., 2022; Mead et al., 2011).

### Spinal Cord Tissue Collection

Mice were anaesthetised with pentobarbitol (30 mg/kg dose; Dolethal), and the chest cavity opened to reveal the heart. The right atrium was severed and 10 ml ice cold 1 × phosphate buffered saline (PBS) was perfused through the left ventricle, followed by 10 ml 4% paraformaldehyde (PFA; Alfa Aesar). The lumbar spinal cord was then dissected and incubated for a further 3–4 hours (h) in 4% PFA before being incubated in sucrose 30% w/v for up to 72 h at 4 °C until sunk. Tissue was then cryo-embedded in OCT compound and stored at − 80 °C. Cryosections were obtained using a Leica CM1860 cryostat at 20 μm thickness and adhered to Superfrost Gold Plus glass slides (VWR). For experiments requiring total internal reflection fluorescence (TIRF) microscopy, 10 μm thick sections were adhered to an Argon Plasma cleaned coverslip (1.0 thickness).

### Immunohistochemistry and aptamer labelling

For immunohistochemistry, slides with spinal cord slices were first heated at 37 °C for 30 min to aid the adherence of the tissue to the glass slides and reduce tissue loss during subsequent wash steps. Slides were washed three times in PBS before being blocked and permeabilised in PBS containing 3% Bovine Serum Albumin (BSA), 1% donkey serum and 0.2% Triton X100 for 2 h at room temperature. Primary antibodies were diluted 1:500 in PBS containing 1.5% BSA and 0.1% Triton X100, and samples were incubated with primary antibody solution for 2 nights at 4 °C. Primary antibodies used include: Anti-VGLUT2 (Mouse, Abcam, ab79157), anti-VGLUT1 (Guinea Pig, Milipore, AB5905), anti-pTDP-43 (Rabbit, ProteinTech, 22309-01). Once incubated, slides are washed five times in PBS over the course of 1–2 h. Secondary antibodies were diluted 1 in 500 in PBS with 0.1% Triton X100, and tissue was incubated in secondary antibody solution for 1.5-2 h at room temperature followed by a further five washes in PBS over the course of 1–2 h. Secondary antibodies used were Donkey-anti-Rabbit conjugated to Alexa Fluor 555 (Abcam, ab150062), Donkey-anti-Mouse conjugated to Alexa Fluor 647 (Abcam, ab150107), and Donkey anti-Guinea Pig conjugated to Alexa Fluor 647 (Jackson Laboratories, 706-605-148). Finally, slides were dried and coverslips of 1.5 thickness were adhered using Prolong Glass Antifade Mountant.

### TDP-43 Aptamer Generation and Labelling

The TDP-43 RNA aptamer conjugated to ATTO 595 was synthesized and purified by HPLC by ATDBio Ltd., as reported previously (Zacco et al., 2022). Spinal cord sections were incubated for at least 1 h with the fluorescently conjugated aptamer prior to imaging and after the primary and secondary antibody incubation steps.

### High-resolution Microscopy

High-resolution confocal-like microscopy was performed using a Zeiss Axio Imager M2 Microscope equipped with an Apotome 2.0 and 63x oil objective lens with a 1.4 numerical aperture (NA), which enables an XY resolution of 320 nm (Broadhead et al., 2020). Illumination was provided by a HXP120 lamp, and images acquired using a digital MRm camera. Exposure times and illumination intensity were kept consistent for analysis within batches of data sets. To map entire hemi-sections of spinal cords to study anatomical diversity between laminae, single optical sections were captured and tiled across half a transverse section of the spinal cord. Images were captured using a single Z-stack plane approximately 3–5 µm depth into the tissue that was kept at the same depth throughout the entire map. Images were stitched together to create whole montage images that were subsequently analysed.

### Airyscan Confocal Microscopy

Sub-confocal resolution images were acquired using a Zeiss LSM800 laser scanning confocal microscope with an Airyscan detector module. The system is based on an Axio ‘Observer 7 microscope. Illumination is provided by a Colibri LED light source, and 405, 488, 561 and 640 nm laser lines. The system is equipped with the Airyscan super-resolution module plus two individual GaAsP PMT detectors. Pixel size was set to 0.04 µm, with a resultant image size of 44.1 × 44.1 µm. Bi-directional scanning was performed with pixel scan time of 2.04 µs and 2x averaging. Multi-channel Z-stack images were acquired using the optimal step-size (0.15 µm) and each channel image was acquired sequentially per Z-plane.

### Gated-Stimulated Emission Depletion Microscopy and Correlative Confocal Microscopy

Multi-channel confocal and gated-stimulated emission depletion (g-STED) microscopy was performed using the Leica SP8 SMD g-STED microscope available at the Edinburgh Super-Resolution Imaging Consortium (ESRIC) hosted by Heriot Watt University. Excitation was provided by a CW super-continuum white light laser source. Depletion was provided by a 594 nm and 775 nm laser. Images were acquired with a 1.4 NA 100× oil STED objective lens and an optical zoom set to provide a resultant image pixel size was 20 nm. Fluorescence was detected using a Leica Hybrid detector, gated at 0.5–8 ns for g-STED images. Confocal and g-STED images were captured sequentially (confocal then g-STED) from regions of interest.

### Aptamer DNA-PAINT

Single molecule localisation microscopy (SMLM) was performed using an Inverted Nikon TI2 microscope (Nikon, Japan) with a 1.49 NA 60x TIRF objective (CFI Apochromat TIRF 60XC Oil, Nikon, Japan) and a ‘perfect focus’ system to autocorrect for z-drift during acquisition. Illumination was provided with 488, 564 and 633 nm laser lines, and detection was provided with an EMCCD camera at 20 Hz (Delta Evolve 512, Photometrics, Tucson, AZ, USA). High-resolution TIRF images were acquired of PSD95-eGFP, VGLUT2 and VGLUT1 by acquiring 100 frames with a 50 ms exposure time and summating the image sequences. Single molecule imaging of aptamer-labelled TDP-43 was performed by acquiring four separate sequences of 1000 frames. Super-resolution images of TDP-43 were generated using Aptamer DNA-PAINT (AD-PAINT) (Whiten et al., 2018). Briefly, the high nanomolar affinity of the aptamer to TDP-43 enables individual binding events to be observed and localized with an accuracy of ∼20 nm. A super-resolution image of TDP-43 is then generated by plotting multiple locations of the aptamer in the tissue.

### Image Analysis

To estimate the size of TDP-43 clusters, the full width at half maximum (FWHM) was calculated from line profiles drawn manually across each punctum. Two lines were drawn perpendicularly across the central point of each punctum, and the average of the two FWHM’s from each line was calculated as the diameter for each TDP-43 cluster.

To quantify the synaptic expression of pTDP-43 from the Airyscan data set, images underwent background subtraction, Gaussian blurring and manual thresholding to segment structures in each image. The Distance Analysis plugin (DiAna) was used to determine the spatial colocalization of synaptic structures and pTDP-43 clusters in 3D (Gilles et al., 2017). As manual thresholding was required, files were renamed and randomised in order to blind the user from experimental condition.

From the SMLM data sets, images were assessed for drift prior to analysis. TDP-43-aptamer images first underwent a background subtraction (rolling ball radius 12 pixels). Using the GDSC SMLM plugin, the Peak Fit function was used to detect single molecule fluorescence events to a Gaussian fit. Localisations were filtered based on a minimum signal strength threshold of 20, a minimum photon count threshold of 50, and a fitting precision threshold of 40 nm. Clusters of TDP-43 from single molecule localisation events were detected using Density-Based Cluster Analysis (DBSCAN) (minimum localisation threshold of 5, clustering distance of 40 nm), and their association with synaptic structures was quantified from their spatial overlap with manually thresholded VGLUT1 and VGLUT2 terminals.

### Statistics

Data handling and preparation of graphs was performed in Microsoft Excel. Statistical tests were performed in Graphpad Prism or SPSS (IBM). Bar charts display error bars denoting the standard error of the mean. Box and whisker plots are used with an inclusive median. Shapiro-Wilk test was used to assess data distribution for normality. One-Way analysis of variance (ANOVA) was performed to quantify inter-regional spinal cord diversity of PSD95 and TDP-43 expression (Figure 2C-E). Analysis of pTDP-43 presence at different synaptic structures and subtypes in healthy control and SOD mice visualised from Airyscan data sets was performed using a multi-factorial ANOVA (Figure 2H,I; Figure 3C,D). Comparisons of pTDP-43 cluster size between healthy control and SOD mice visualised from Airyscan data sets (across n=5-6 mice) was performed using a Two-sample T-Test (Figure 3E,F). Comparisons of pTDP-43 cluster size between healthy control and SOD mice (n=100 clusters) visualised from g-STED data set was performed using a Mann-Whitney U Test (Figure 3H).

**Figure 1.**
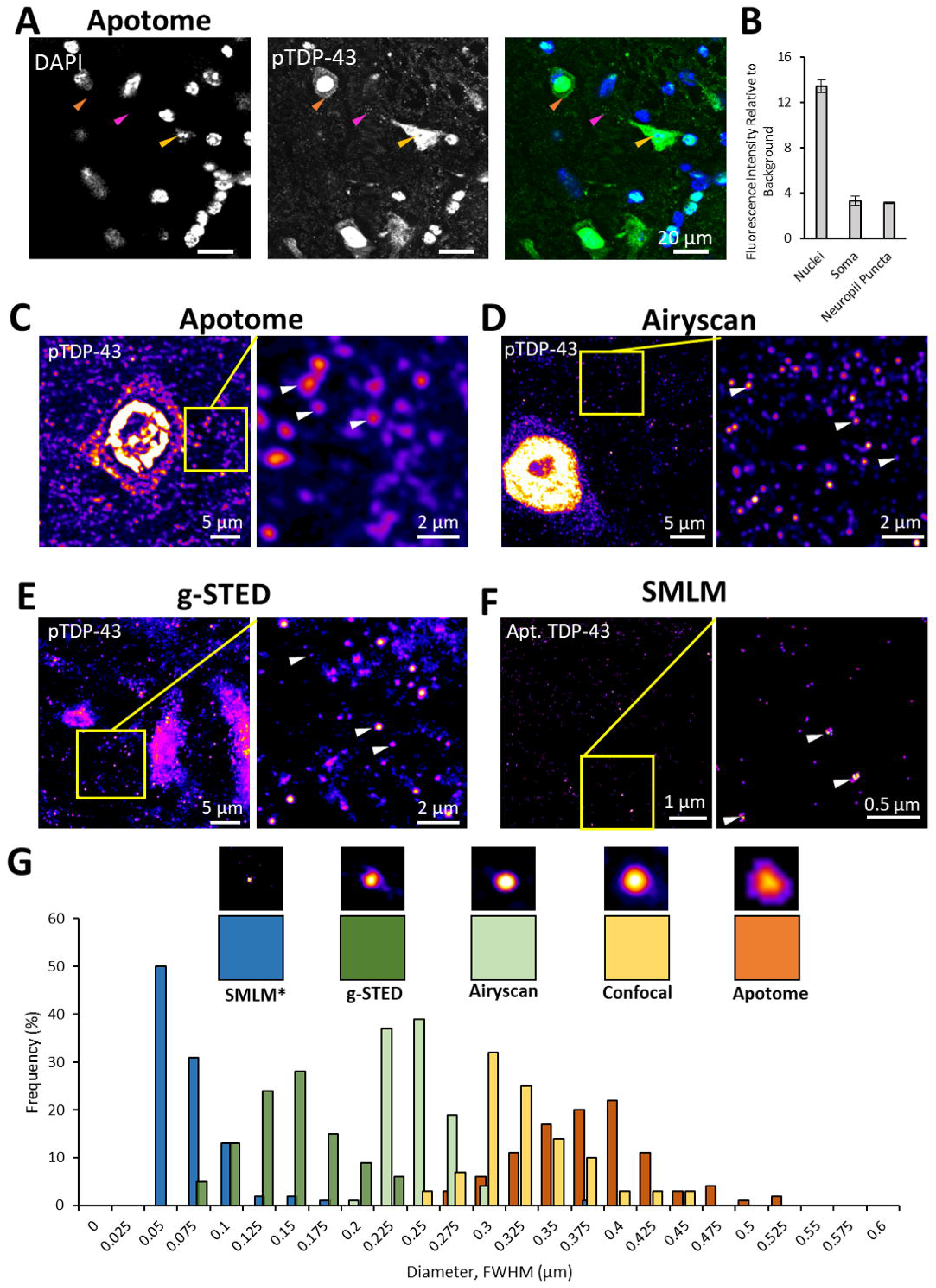
Visualising TDP-43 Clusters in the Mouse Spinal Cord Using Super-Resolution Microscopy. This figure demonstrates pTDP-43 and TDP-43 expression in the mouse spinal cord visualised using a range of different microscopy techniques. A. A high-resolution Apotome microscope was used to visualise pTDP-43 expression (green) in the ventral horn of the mouse lumbar spinal cord, co-stained with the nuclear marker, DAPI (blue). The images indicate nuclear expression (yellow arrow) and somatic expression (orange arrow) and clusters in the neuropil (magenta arrow). B. The intensity of pTDP-43 expression in nuclei, apparent somatic and neuropil clusters were measured compared to the background intensity. C-E. Example images of pTDP-43 labelling are shown using different microscopes with different resolutions, including the Zeiss Apotome (C), Zeiss Airyscan (D) and Leica g-STED (E). White arrows indicate some example pTDP-43 clusters. F. Example images of TDP-43 labelled using a fluorescently conjugated aptamer and visualised using AD-PAINT. G. The sizes of apparent pTDP-43 clusters and aptamer labelled-TDP-43 clusters (*) as measured from the FWHM of line profiles from across clusters are plotted in a frequency histogram for each microscopy method. Example images of TDP-43 clusters visualised with each method are displayed in 1 µm x 1 µm boxes with their colour coded below.

**Figure 2.**
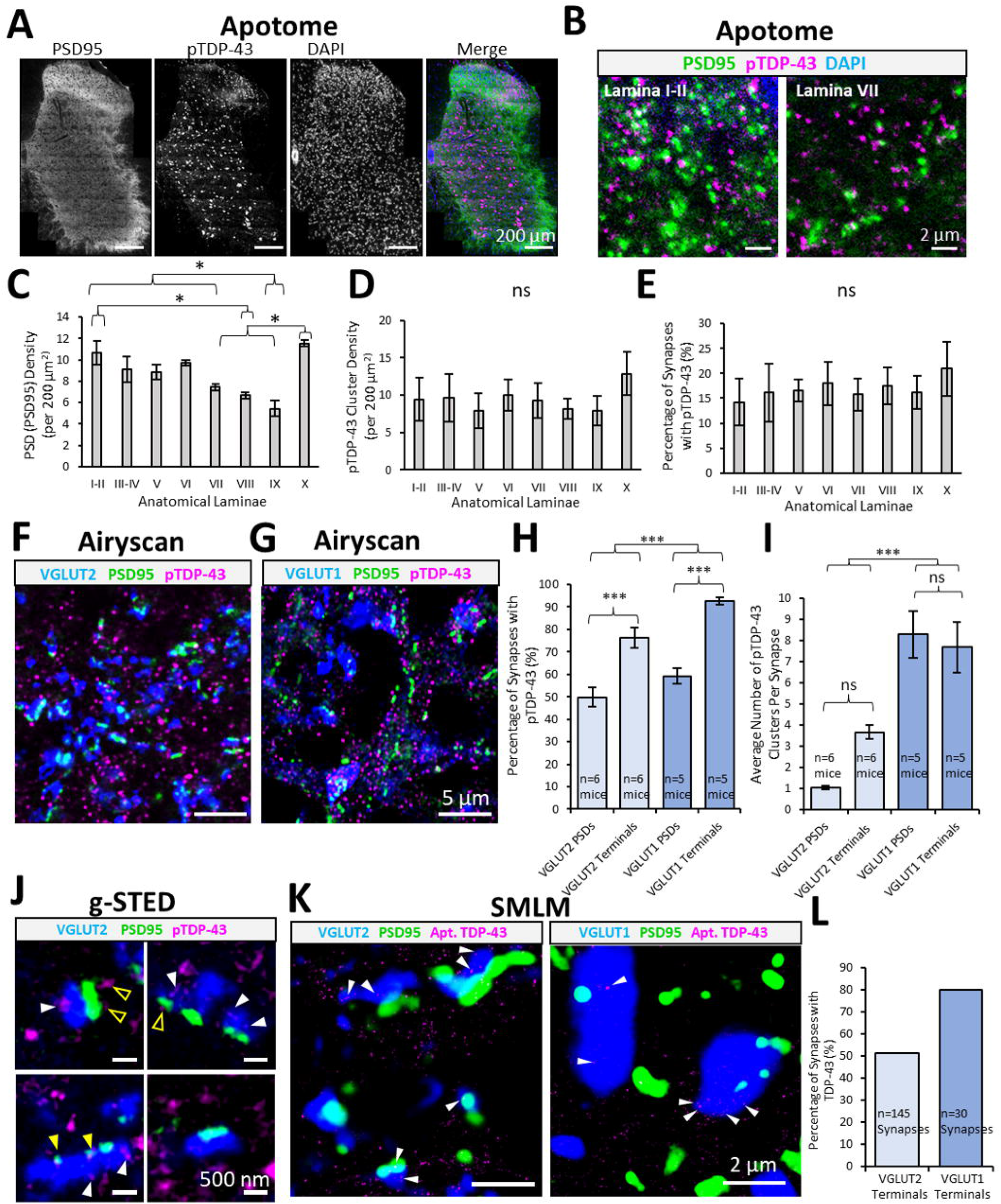
Quantifying the Synaptic Expression of TDP-43 in the Mouse Spinal cord. The synaptic expression of TDP-43 in the mouse lumbar spinal cord was characterised using a range of high-resolution and super-resolution microscopy methods. A. The synaptic expression of pTDP-43 clusters was mapped throughout different anatomical laminae of the lumbar mouse spinal cord using a high-resolution tiling approach (Apotome). Synapses were labelled genetically with eGFP-tagged PSD95 (green), and nuclei were labelled with DAPI (blue). B. Example images of pTDP-43 clusters and PSD95-eGFP puncta in the dorsal laminae I-II and ventral laminae VII, demonstrating synaptic diversity between sensory and motor regions of the spinal cord. C. Graph plots the synaptic density (PSD95-eGFP puncta per unit area) across different anatomical laminae, demonstrating significant inter-regional diversity in the distribution of synapses. D. Graph displaying no difference in the density of pTDP-43 clusters between different anatomical subregions. E. Graph displaying no difference in the percentage of PSD95 synapses expressing pTDP-43 clusters between different anatomical subregions. F-G. To ask whether there was diversity or specificity in the synaptic localisation of pTDP-43 in different synapse structures and subtypes, the expression of pTDP-43 clusters was visualised in greater detail using the Zeiss Airyscan to quantify postsynaptic expression (by 17olocalization with PSD95-eGFP), and presynaptic expression (by 17olocalization with VGLUT2 terminals (F) or VGLUT1 terminals (G)). Images were acquired from ventral horn lamina IX. H. Graph plotting the percentage of VGLUT2-associated PSDs and terminals (lighter blue bars) and VGLUT1-associated PSDs and terminals (darker blue bars) containing pTDP-43 clusters. I. Graph plotting the average number of pTDP-43 clusters per VGLUT2-associated PSDs and terminals (lighter blue bars) and VGLUT1-associated PSDs and terminals (darker blue bars). J. Three-colour g-STED super-resolution images further characterise the subsynaptic distribution of pTDP-43 expression in VGLUT2-associated synapses. Many pTDP-43 clusters were observed presynaptically (white filled arrows), with some cases of pTDP-43 clustering within the PSD (yellow filled arrows) and other cases of clusters located nearby – but not directly at – the PSD (yellow outlined arrows). K. AD-PAINT was used to visualise aptamer-labelled TDP-43 in PSD95-eGFP mouse spinal cord sections immunolabelled with either VGLUT2 or VGLUT1 to quantify the synaptic diversity of TDP-43 expression. J. Graph plotting the percentage of VGLUT2 and VGLUT1 presynaptic terminals expressing aptamer-labelled TDP-43 clusters, confirming that TDP-43 appeared to be more frequently expressed in VGLUT1 terminals than VGLUT2 terminals.

**Figure 3.**
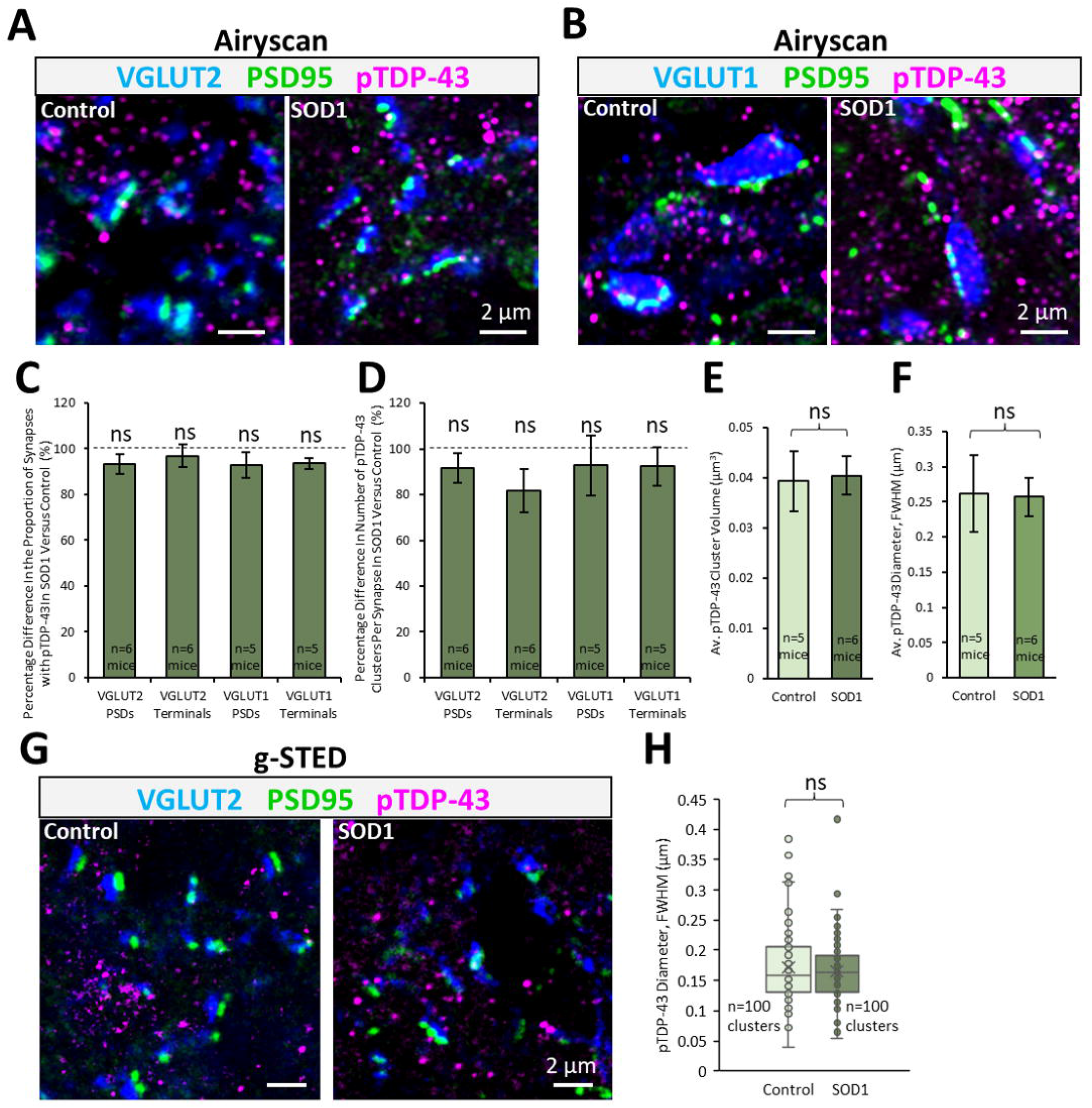
Synaptic pTDP-43 clusters are no different between healthy control and SOD1 mice. To investigate whether the ALS mouse model, SOD1, displays any change in the synaptic expression or pTDP-43, excitatory synapses and pTDP-43 clusters were visualised in symptomatic stage SOD1 mice (112 days old) and age matched healthy controls. A-B. Mouse spinal cord sections expressing PSD95-eGFP were co-labelled for pTDP-43 and either VGLUT2 (A) or VGLUT1 (B) to visualise PSDs, and presynaptic terminals from different synapse subtypes. 3D Z-stack acquisitions were captured using the Airyscan microscopy. C. Graph showing no difference in the percentage of synapse subtypes expressing pTDP-43 clusters in SOD1 mice compared to controls. D. Graph showing no difference (as a percentage compared to controls) in the number of pTDP-43 clusters within different synapse subtypes in SOD1 mice. E. The size of synaptically expressed pTDP-43 clusters, as measured from the volume of the thresholded cluster, was no different between control and SOD1 mice. F. The size of the synaptically expressed pTDP-43 clusters, as measured from the FWHM, was no different between control and SOD1 mice. G. g-STED microscopy images were captured of pTDP-43 clusters in VGLUT2-associated synapses in a healthy control and SOD1 mouse. H. The size of the synaptic pTDP-43 clusters, as measured from the FWHM, was no different between control and SOD1 mice.

## Results

### Characterisation of TDP-43 Expression in Mouse Spinal Cord Neuropil

Immunolabelling of pTDP-43 was performed in upper lumbar spinal cord sections obtained from adult mice. Images were captured using a range of fluorescence microscopes, each offering different spatial resolutions: the Zeiss Apotome (∼320 nm), Zeiss Airyscan (∼200 nm) and Leica g-STED (∼75 nm). In addition, single-molecule localisation microscopy (SMLM) was employed to visualise TDP-43 using an aptamer based labelling approach (resolution ∼ 20 nm) (Zacco et al., 2022).

Using the high-resolution Zeiss Apotome microscope, pTDP-43 was found predominantly at the nucleus of cells but there was also considerable punctate labelling in cell bodies and in the neuropil (Fig. 1 A). The fluorescence intensity of nuclear pTDP-43 expression was 12-fold higher than the background, while puncta in the nearby soma and neuropil were 3-fold brighter than the background intensity (Fig. 1B).

In order to investigate the size and morphology of TDP-43 clusters, the ventral horn of adult mouse spinal cords immunolabelled with pTDP-43 were visualised using a Zeiss Apotome, Zeiss Airyscan, Leica Confocal and g-STED microscope (Fig. 1C-F). The average diameter of TDP-43 clusters, calculated from the FWHM of line profiles of 100 randomly selected puncta, visualised using each method, is plotted in the histogram in panel Fig. 1G. pTDP-43 clusters visualised with the Zeiss Apotome measured 366 ± 51 nm (mean ± standard deviation) (Fig. 1C,G). Clusters measured with the Airyscan measured 236 ± 20 nm (Fig. 1D,G). Clusters of pTDP-43 visualised using confocal microscopy with the Leica Sp8 measured 316 ± 42 nm (Fig. 1G). The same clusters were then visualised and analysed using STED acquisition settings, and measured 135 ± 36 nm (Fig. 1E,G). In separate acquisitions, using gating in addition to the STED (g-STED) improved the resolution slightly resulting in pTDP-43 clusters measuring 115 ± 52 nm diameter (n=21 pTDP-43 clusters) (SI. Fig 1A). However, the combination of gating and high STED depletion powers resulted in a considerably reduced signal to noise ratio, and thus many pTDP-43 clusters that could be detected with confocal microscopy were no longer detectable with g-STED (SI. Fig. 1B). The difference in apparent size of pTDP-43 clusters using the different methods demonstrates the resolution capabilities of each system.

An alternative approach to immunolabelling is to use DNA or RNA-based aptamers, designed to bind to key peptides or protein sequences of interest. We employed an in silico-designed aptamer that has recently been demonstrated to specifically target TDP-43 clusters and aggregates *in vitro* (Zacco et al., 2022). Mouse spinal cord sections were incubated in the fluorescently conjugated TDP-43 aptamer solution, and super-resolution images were generated using Aptamer DNA-PAINT (AD-PAINT) (Whiten et al., 2018). Clusters of aptamer-labelled TDP-43 measured 62 ± 39 nm using FWHM analysis from line intensity profiles, supporting the concept of small localised clusters of TDP-43 in the neuropil (Fig. 1F-G). The resolution of this approach tested across 3 representative images calculated as 37.8 ± 0.08 nm, as measured by Fourier Ring Correlation (Nieuwenhuizen et al., 2013).

### Synaptic Expression of TDP-43

We next determined whether the TDP-43 clusters observed in the neuropil were associated with synapses by visualising pTDP-43 along with synaptic markers using high-resolution and super-resolution microscopy in healthy adult mouse lumbar spinal cords. The genetically encoded PSD95-eGFP marker was used to detect the PSDs of excitatory synapses.

Using the high-resolution Zeiss Apotome, excitatory synapses were mapped throughout different anatomical laminae of the mouse lumbar spinal cord (laminae I-II, III-IV, V, VI, VII, VIII, IX and X), and the number of synapses and pTDP-43 clusters was quantified (Fig. 2A-E). Similar to previous observations (Broadhead et al., 2020), synaptic mapping in mouse spinal cords showed significant diversity between anatomical laminae in the number of excitatory synapses per unit area (PSD density) (Fig. 2C; n=4 mice, F(7)=8.7, p<0.0001). This diversity was characterised by a much greater number of synapses in the dorsal laminae compared to the ventral laminae (Fig 1B-C). In contrast, there was no difference in the density of pTDP-43 clusters between different anatomical laminae (Fig. 2D; F(7)=0.4, p=0.868). Nor was there any difference in the proportion of PSD95 synapses that colocalised with pTDP-43 clusters between different laminae (Fig. 2E; F(7)=0.2, p=0.979).

To accurately quantitate the presynaptic and postsynaptic expression of pTDP-43 clusters at excitatory synapses, PSD95-eGFP mouse spinal cord sections were immunolabelled with pTDP-43 along with either anti-VGLUT2 (Fig. 2F) or anti-VGLUT1 (Fig. 2G). In the ventral horn of the mammalian lumbar spinal cord, VGLUT1 terminals arise from corticospinal pathways and mechanoreceptive terminals from sensory neurons within the dorsal root ganglia (Ni et al., 2014; Todd et al., 2003). Meanwhile, the majority of VGLUT2 terminals derive from local spinal neurons, with a smaller number corresponding to nociceptive inputs and descending inputs from the rubrospinal and vestibulospinal tracts (Du Beau et al., 2012; Lagerström et al., 2010; Todd et al., 2003). 3-D z-stacks were acquired in the ventral horn laminae IX – where the lateral motor neuron pools reside – and the expression of pTDP-43 clusters inside populations of pre- and post-synaptic structures were analysed (Fig. 2H-I).

From analysis of pTDP-43 clusters in VGLUT2-associated synapses (n=6 mice), pTDP-43 clusters were identified at 49 ± 9 % of VGLUT2-associated PSDs compared to 76 ± 10 % of VGLUT2 presynaptic terminals (Fig. 2H). From analysis of pTDP-43 clusters in VGLUT1-associated synapses (n=5 mice), a similar proportion of PSDs contained pTDP-43 clusters compared to VGLUT2-associated PSDs (55 ± 7 %), while significantly more VGLUT1-associated terminals contained pTDP-43 clusters (87 ± 5 %). Overall, pTDP-43 clusters were more frequently expressed at VGLUT1-associated synapses than VGLUT2-associated synapses (F(1)=23.1, p<0.0001), and more frequently associated with presynaptic terminals than PSD’s (F(1)=143.7, p<0.0001). Similarly, when the number of pTDP-43 puncta per synapse was analysed (Fig. 2I), it was observed that VGLUT1 synapses had a much greater number of pTDP-43 clusters per synapse than VGLUT2 synapses (F(1)=91.6, p<0.0001), though there was no difference in the number of pTDP-43 clusters between presynaptic terminals and PSDs (F(1)=0.03, p=0.858).

Using g-STED microscopy, the subsynaptic expression of pTDP-43 was also elucidated (Fig. 2J). VGLUT2 and PSD95 positive excitatory synapses displayed pTDP-43 clusters predominantly pre-synaptically. Some cases of pTDP-43 cluster colocalization with PSD95 were observed, while in other cases pTDP-43 clusters resided >100 nm from the PSD, suggesting that some clusters may reside within the dendritic spine head or shaft, but away from the PSD itself.

In addition, AD-PAINT was used to investigate whether aptamer-labelled TDP-43 displayed a differential distribution in synapse subtypes. PSD95-eGFP, VGLUT1 and VGLUT2 were immunolabelled in mouse spinal cord sections along with the incubation of the anti-TDP-43 aptamer and images were captured (across 1 or 2 sections from the same mouse). From analysis of 145 VGLUT2 synapses and 30 VGLUT1 synapses, we observed that 51% of VGLUT2 terminals expressed TDP-43 clusters compared to 80% of VGLUT1 terminals.

These data demonstrate that TDP-43 is expressed and localised in clusters at synapses, with a more prominent presynaptic distribution, and a heterogenous expression between different subtypes of synapses.

### Synaptic Expression of TDP-43 in a Mouse Model of ALS

Various reports indicate a key role for TDP-43 pathology in synaptic dysfunction in ALS. The ALS mouse model, SOD1^G93a^ (herein referred to as SOD1) displays many cellular and behavioural phenotypes that recapitulate the human disease, including synaptic dysfunction and degeneration. Although many reports indicate that the SOD1 mouse model displays no TDP-43-associated pathology, the synaptic presence of TDP-43 in this model has not been investigated using nanoscopy techniques that may detect more subtle changes. We therefore assessed the subsynaptic expression of pTDP-43 in populations of synapses in the lamina IX motor pools of upper lumbar spinal cords from both healthy control and SOD1 mice. The SOD1 mice were cross-bred with PSD95-eGFP mice such that all progeny were heterozygous for the PSD95-eGFP mutation, with half the progeny expressing the SOD1 mutation, and the other half used as control mice. Mice were aged 16 weeks old (112-days); a time-point we have previously demonstrated to display significant synapse loss (Broadhead et al., 2022). Spinal cords were immunolabelled for pTDP-43 and either VGLUT2 (Fig. 3A) or VGLUT1 (Fig. 3B), and 3D images were acquired using the Zeiss Airyscan.

Our examination of the proportion of different synaptic structures (VGLUT2-associated PSDs and terminals, and VGLUT1-associated PSDs and terminals) containing pTDP-43 clusters revealed no differences between controls and SOD1 mice (Fig. 3C; F(1)=2.5, p=0.123). There was also no difference in the number of pTDP-43 clusters per synapse between controls and SOD1 mice (Fig. 3D; F(1)=0.8, p=0.374). Analysis of all pTDP-43 clusters expressed at either VGLUT1 or VGLUT2 synapses, demonstrated that there was no difference in their size measured from the volume of each cluster (Fig. 3E; T(f)=0.01, p=0.991). Using a more accurate method to analyse the size of pTDP-43 clusters from a smaller subset of VGLUT2 synapses, diameter estimated from the FWHM of line profiles of pTDP-43 clusters, we again found no significant difference in the size of synaptic pTDP-43 clusters between control and SOD1 mice (Fig. 3F, t(9)=0.18, p=0.863). Using g-STED microscopy to visualise pTDP-43 clusters in VGLUT2-associated synapses from a single healthy control and single SOD1 mouse, it was further confirmed that synaptic pTDP-43 clusters were no different in size between SOD1 mice and healthy controls (Fig. 3G-H; U=4798, p=0.622).

## Discussion

### Nanoscale Synaptic Clustering of TDP-43

Our study reveals nanoscale synaptic expression of TDP-43 in the ventral horn of the lumbar spinal cord in adult mice. This was apparent using both an anti-pTDP-43 antibody and an anti-TDP-43 aptamer, visualised with a range of high-resolution and super-resolution microscopy techniques. While there is considerable literature on the role of TDP-43 in synaptic function in health and disease (Bak et al., 2022; Dyer et al., 2021; Fogarty et al., 2016; Godena et al., 2011; Jiang et al., 2019), our findings offer substantial subsynaptic characterisation of TDP-43 that support the conclusion that TDP-43 may have direct implications on synaptic function. The clusters of TDP-43 at synapses appear to be diffraction limited in size – measuring 60nm in diameter using AD-PAINT, or just over 100 nm in diameter using g-STED microscopy. These measurements are largely consistent with other quantitative assessments of TDP-43 clusters using similar super-resolution microscopy techniques and electron microscopy in mammalian cell lines and tissue (Johnson et al., 2009; Kao et al., 2015; Wong et al., 2022; Zacco et al., 2022). Interestingly, recent findings using super-resolution microscopy discerned that larger inclusions of TDP-43, not smaller ones, were more strongly associated with neuronal toxicity (Cascella et al., 2022). Nevertheless, super-resolution techniques offer the potential for capturing nanoscale changes that may precede or predict the formation of larger TDP-43 aggregates and inclusions.

The apparent size of TDP-43 clusters measured is highly dependent on the resolution of the system. Furthermore, aptamer labels are much smaller than primary-secondary antibody complexes. Therefore, the spatial localisation the aptamer-conjugated fluorophore to the protein target of interest is more accurate, while antibody approaches may displace the signal from the target protein, resulting in seemingly larger structures. However, our data sets revealed striking homogeneity in the distribution of TDP-43 cluster sizes using each methodology, suggesting that synaptic TDP-43 clusters in the mouse spinal cord are relatively similar in size and shape. Each method incurs its own advantages and limitations. While AD-PAINT provides optimal resolution of TDP-43 clusters, it incurs slow acquisitions, generating large data sets that require considerable computational power (>1Gb per acquisition), that are complex and time-consuming to process and analyse. The Apotome system is remarkably fast and facilitates high-resolution imaging across larger areas of tissue, but its resolution is less than a conventional confocal microscope. Both the Airyscan and g-STED methods offered reasonable improvements in resolution beyond the diffraction limit of light (though inferior to SMLM), and are relatively ‘user friendly’ for non-specialist microscopists in terms of sample preparation, data set size and analysis.

Nanoscale clusters of TDP-43 were found in both VGLUT1- and VGLUT2-associated presynaptic terminals in regions where motor neurons reside. The more frequent and predominant expression of TDP-43 clusters in VGLUT1-associated synapses may suggest a more prominent role for TDP-43 in mRNA transport or protein translation at these subtypes of synapses, which may correspond to primary mechanoreceptive terminals or corticospinal pathways (Brown & Fyffe, 1978; Ni et al., 2014; Schultz et al., 2017; Todd et al., 2003). The differential expression of TDP-43 between the two subtypes of synapses, however, could incur selective vulnerability of certain synapse subtypes to TDP-43 related neuropathologies, as is observed in ALS. While TDP-43 appeared to be more predominantly presynaptically expressed, our data also indicate the presence of postsynaptic TDP-43 clusters by its partial colocalization with PSD95, which has also been observed in other studies using PSD95 as a postsynaptic marker (Henstridge et al., 2018).

Our data do not reveal any significant difference in the cluster size or synaptic expression of pTDP-43 in SOD1 mice compared to healthy littermate controls. Previous reports have provided conflicting results regarding the presence or timescale of TDP-43 pathology in the SOD1^G93a^ mouse model. Shan et al., (2009) demonstrated cytoplasmic inclusions of TDP-43 in ‘end-stage’ mice that were approximately 170 days old probed with an anti-TDP-43 antibody. Jeon et al., (2019), demonstrated increased immunoreactivity for pTDP-43 and the C-terminal fragment of TDP-43, but not the N-terminal of TDP-43, in SOD1 mice aged 120 days – the same age used in our study. However, other studies have demonstrated no such TDP-43 pathology at similar early symptomatic stage mice as well as sporadic ALS patients carrying SOD1 mutations (Henstridge et al., 2018; Robertson et al., 2007; Turner et al., 2008). It could be that TDP-43 pathology and synaptopathy in SOD1 mice occur through independent mechanisms, despite the fact that TDP-43 is present at synapses and that other findings from human post-mortem studies suggest a correlation between TDP-43 pathology and synaptic loss (Henstridge et al., 2018). Similarly, other synapse subtypes received by motor neurons are also subject to changes, including inhibitory glycinergic synapses (Allodi et al., 2021) and modulatory cholinergic synapses (Bak et al., 2022; Herron & Miles, 2012; Witts et al., 2014). The approaches used in this study would be highly suitable for identifying nanoscopic TDP-43-associated synaptopathy in other such synapse subtypes in ALS.

Mutations in a range of different genes, including most commonly C9ORF72, TDP43, FUS and SOD1, can all lead to some form of motor neuron disorder or the related cognitive disorder frontotemporal dementia. It is possible that the synapse may act as a key subcellular locus of convergence for these multiple different disease pathways. For example, C9orf72 has been shown to interact with the presynaptic protein, Synapsin-I, and it’s perturbed interactions with Synapsin-I in ALS may be associated with excitatory synapse dysfunction and loss (Bauer et al., 2022; Devlin et al., 2015; Perkins et al., 2021). Super-resolution microscopy techniques have been used to identify the presynaptic localisation of FUS at excitatory synapses, and mutations in FUS are similarly implicated in excitatory synapse dysfunction and loss (Deshpande et al., 2019; Sahadevan et al., 2021; Schoen et al., 2015). There is extensive literature on synaptic dysfunction associated with mutations in the SOD1 gene (Allodi et al., 2021; Avossa et al., 2006; Bączyk et al., 2020; Broadhead et al., 2022; Fogarty, 2019; Fogarty et al., 2015; Herron & Miles, 2012; Naumenko et al., 2011; Rei et al., 2020). Finally, structural and functional synaptic changes have been demonstrated in animal models with TDP-43 mutations (Bak et al., 2022; Dyer et al., 2021; Heyburn & Moussa, 2016; Jiang et al., 2019) and human iPSC neurons from patients with TDP-43 mutations (Devlin et al., 2015), while our findings, along with other published results, support synaptic roles for TDP-43 based on its presynaptic and postsynaptic expression (Johnson et al., 2009; Kao et al., 2015; Wong et al., 2022; Zacco et al., 2022). There is also evidence that misfolding of TDP-43 and FUS proteins may induce pathological misfolding of SOD1 through prion-like mechanisms (Pokrishevsky et al., 2016). The small confined domain of the synapse may increase the chance of protein interaction inducing misfolding, as well as facilitate the prion-like spread of proteins between neurons using exosome-dependent mechanisms (Shimonaka et al., 2016; Westergard et al., 2016), thus further supporting the notion that the synapse could be a highly vulnerable common pathway for different ALS/FTD disease mechanisms.

In summary, our quantitative super-resolution imaging provide novel characterisation of the subcellular distribution of TDP-43 and nanoscale organisation of protein clusters. These techniques are valuable for understanding the role of TDP-43 in neuronal function. The sensitivity and resolution of such techniques will be valuable in identifying subtle early-stage aggregations of TDP-43, as it is known that complex aggregates form during early pathogenesis of ALS (Chen & Cohen, 2019; Ling, 2018; Ling et al., 2013). Such information will help advance our understanding of early stage biomarkers of disease and reveal novel therapeutic avenues that target synaptopathy.

## Acknowledgements

We would like to acknowledge the following funders: Motor Neurone Disease (MND) Association UK (Miles/Apr18/863-791), Chief Scientist Office, RS Macdonald Charitable Trust, the European Research Council (ERC) under the European Union’s Horizon 2020 Research and Innovation Programme (695568 SYNNOVATE), Simons Foundation Autism Research Initiative (529085), and the Wellcome Trust (Technology Development grant 202932).

