## Supplementary Figures and Tables for "Synaptic Expression of TAR-DNA-Binding Protein 43 in the Mouse Spinal Cord Determined Using Super-Resolution Microscopy"

**Supplementary Figure.**

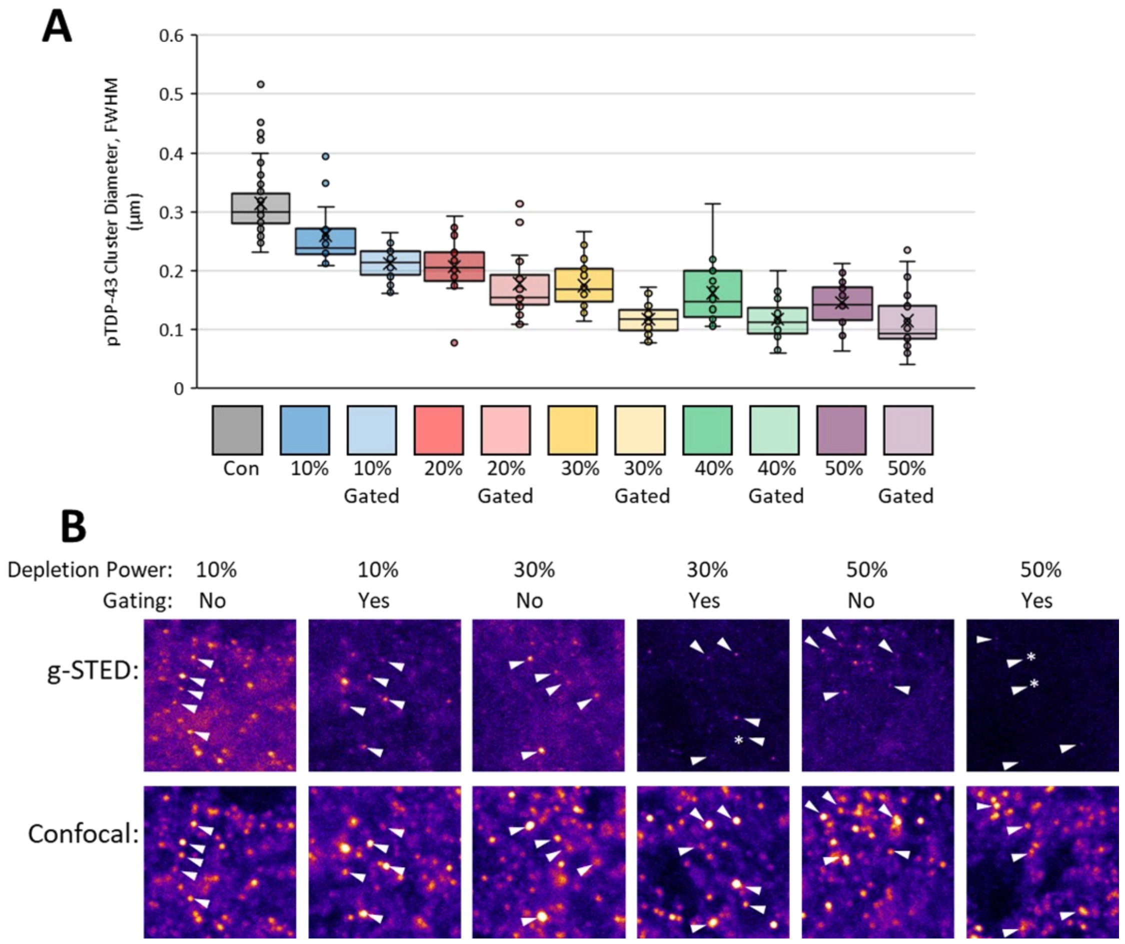

**SI. Figure 1. Quantitative and qualitative assessment of gated STED microscopy imaging settings for visualising pTDP-43 clusters.**

Images were captured of pTDP-43 clusters in the ventral horn of the upper lumbar mouse spinal cord. The 775nm depletion laser power, and use of gating were titrated, and the size of pTDP-43 clusters were measured from the FWHM of intensity line profiles. A. Increasing STED laser power improved the resolution, as shown by the reduced size of pTDP-43 clusters. The presence of gating also improved resolution with respect to each depletion laser power. B. pTDP-43 clusters were still visible in confocal and g-STED conditions (white arrows). However, at 50% depletion laser power, and the use of gating, some TDP-43 clusters were no longer detectable in g-STED despite their presence in confocal imaging (stars), suggesting reduced signal-to-noise-ratio in the acquisitions.
